# Repurposing anti-phage defenses to differentially arrest the viral lifecycle reveals the regulatory logic of a parasitic satellite

**DOI:** 10.64898/2026.04.03.716414

**Authors:** S. Tansu Bagdatli, Kimberley D. Seed

## Abstract

Mobile genetic elements frequently encode defense mechanisms to protect their bacterial hosts from viral attack. In *Vibrio cholerae,* these defensive elements include phage-inducible chromosomal island-like elements (PLEs), which are phage satellites that act as highly specialized parasites of the lytic phage ICP1. While PLE transcriptional activation upon ICP1 infection is known to be temporally regulated, the underlying regulatory logic and dependencies on the progression of the phage’s developmental program required for activation remain unclear. In this study, we took a novel approach to define these dependencies by introducing independent anti-phage defense systems, BREX and DarTG, as molecular roadblocks to impede the ICP1 lifecycle. We discovered that, for both ICP1 and PLE, late-stage gene expression is fundamentally uncoupled from genome replication, representing a striking departure from the standard paradigm for double-stranded DNA phages. While BREX restricts ICP1 to an immediate-early transcriptional state that stalls PLE activation, DarTG allows the phage to execute its full transcriptional cascade despite the total block in DNA replication. This permissive environment provides the necessary cues for complete PLE induction, revealing that the extent of ICP1 transcriptional progression is a key determinant of PLE transcriptional activation. Unlike other phage satellites that rely on a single cue for activation, our results demonstrate that PLE uses a progressive licensing strategy that relies on multiple cues tied to milestones in the phage’s developmental program. This regulatory architecture ensures robust PLE activation resilient to phage escape.

## INTRODUCTION

Phages, the viruses that infect bacteria, shape bacterial population dynamics by infecting, manipulating, and killing their bacterial hosts. This dynamic interaction exerts intense selective pressure, driving bacteria to evolve a diverse arsenal of anti-phage defense systems ^1^. These systems are frequently encoded by mobile genetic elements (MGEs) ^2^ and can be broadly categorized by the timing and underlying mechanism of their action. Early-acting, cell-autonomous systems, such as restriction modification systems and bacteriophage exclusion (BREX), distinguish self from non-self DNA, often preventing phage DNA replication immediately upon entry ^3–5^. In contrast, abortive infection (Abi) systems may be triggered later and function as a communal defense, in which the infected cell dies upon phage detection, preventing phage proliferation and safeguarding the surrounding bacterial population ^2,6^. The DarTG toxin-antitoxin system exemplifies this strategy, whereby the DarT toxin chemically modifies phage DNA ^7^ to halt its replication, triggering cell death that protects uninfected neighbors ^8^.

While systems like DarTG are standalone defenses, some MGEs have evolved to function as highly specialized Abi systems that parasitize invading phages. In *Vibrio cholerae*, the agent of cholera, these elements are a distinct family of phage satellites known as <u>p</u>hage-inducible chromosomal island-like <u>e</u>lements (PLEs). *V. cholerae* is persistently attacked by the lytic myovirus ICP1 during human infection ^9,10^, and PLEs provide highly specific defense against ICP1 by acting as both selfish phage parasites and effective abortive infection systems ^10,11^. The PLE life cycle is tightly coupled to the ICP1 life cycle. Upon ICP1 infection, PLE is transcriptionally activated and executes a temporally regulated gene expression cascade comprising early, middle, and late phases that mirrors the phage’s own rapid 20-minute infection cycle ^11,12^. During this time, PLE excises from the *V. cholerae* chromosome ^13^, replicates to high copy numbers ^14^, and hijacks ICP1 structural proteins to package its own genome into PLE transducing particles ^15^, while simultaneously abolishing progeny production by blocking late-stage phage morphogenesis ^15–17^.

As phage satellites, PLEs belong to a widespread class of MGEs defined by their parasitism of a helper phage to promote their own mobilization ^18^. The canonical model for how phage satellites sense and respond to their helper is based on the *Staphylococcus aureus* pathogenicity islands (SaPIs). In the absence of helper phage, SaPIs are maintained in a dormant state by a master repressor; upon infection, a single phage protein acts as an anti-repressor, triggering satellite activation ^19,20^. However, PLE activation appears to deviate from this canonical single-trigger model. Unlike SaPI helper phages, which can escape satellite induction via point mutations in the non-essential anti-repressor ^20^, spontaneous ICP1 mutants that can escape PLE-mediated restriction have not been reported. This absence of escape mutants suggests that PLE likely responds to an essential phage protein that is intolerant of mutation, or that it relies on multiple phage proteins to trigger its temporal gene expression cascade. While we have previously established that the bacterial protein H-NS silences the model variant PLE1 in the absence of infection ^21^, the precise viral cues that trigger PLE activation remain unknown.

Given the absence of an apparent single SaPI-like activation trigger, we hypothesized that the transcriptional regulation of PLE is coupled to the progression of the ICP1 life cycle. To interrogate these dependencies, we used BREX and DarTG defense systems as distinct molecular tools to limit the progression of ICP1’s life cycle. Both systems block ICP1 genome replication ^22,23^, but they represent distinct early-acting and abortive strategies. Critically, it was unknown how these different defenses impact the progression of the phage’s temporal transcriptional cascade. We reasoned that despite their shared inhibition of phage genome replication, these systems might serve as temporal roadblocks at different stages of the phage’s developmental program. By characterizing these barriers, we aimed to decouple phage replication from transcriptional progression. This approach allowed us to determine whether PLE activation is triggered by initial phage gene expression or requires specific milestones in the phage’s developmental program, providing critical insight into whether this satellite relies on a single trigger or multiple, temporally distributed cues.

By employing this comparative approach, we uncovered a critical divergence in how BREX and DarTG defenses impact phage development. While both block ICP1 genome replication ^22,23^, we found they impose drastically different constraints on phage gene expression. In the presence of BREX, phage gene expression is potently repressed. In contrast, DarTG—despite blocking replication—allows the phage to progress to late-stage gene expression. This finding demonstrates that for ICP1, the activation of late genes encoding structural and lysis proteins occurs independently of successful genome replication—a striking deviation from the standard paradigm for double-stranded DNA viruses, including phages ^24,25^. We leveraged these contrasting transcriptional landscapes to probe the requirements for PLE activation and found that PLE gene expression is strictly dependent on the progression of the phage’s transcriptional program. In the presence of BREX, PLE gene expression is severely attenuated and varies by operon, whereas in the DarTG background, PLE executes its full transcriptional activation program. Notably, this reveals that PLE mirrors the phage’s capacity to uncouple transcriptional progression from genome replication, as the satellite activates late gene expression even when its genome replication is blocked. Collectively, our findings support a model in which PLE activation is not triggered by a single factor but is intimately tied to the global transcriptional state of its helper phage.

## RESULTS

### Comparative transcriptomics reveals distinct global impacts of BREX and DarTG on PLE and phage gene expression

We previously identified two anti-phage defense systems that block the initiation of ICP1 genome replication in clinical isolates of *V. cholerae*: BREX, encoded by an integrative and conjugative element ^22^, and DarTG, a toxin-antitoxin system encoded by a distinct phage defense element ^23^. To enable appropriate comparisons, we separately introduced the genetic elements encoding BREX and DarTG into the same parental *V. cholerae* strain harboring the most well-studied PLE variant, PLE1 (hereafter PLE) (Fig. 1A). We then generated isogenic controls by deleting the genes responsible for inhibiting ICP1 replication within each element (see methods). This process yielded two matched sets of strains: functional defense (BREX(+) and DarTG(+)) and replication-permissive controls (BREX(-) and DarTG(-)). This design allows us to compare infection restricted by distinct defenses, both of which inhibit ICP1 genome replication (Fig. S1A), with one in which the phage developmental program proceeds. Although PLE ultimately provides abortive defense by blocking late-stage phage assembly ^15–17^, it exerts minimal impact on ICP1 genome replication and transcription ^12,14^. Consequently, the defense-deficient strains serve as replication-permissive controls, allowing us to assess the specific transcriptional constraints imposed by BREX and DarTG.

**Fig. 1:**
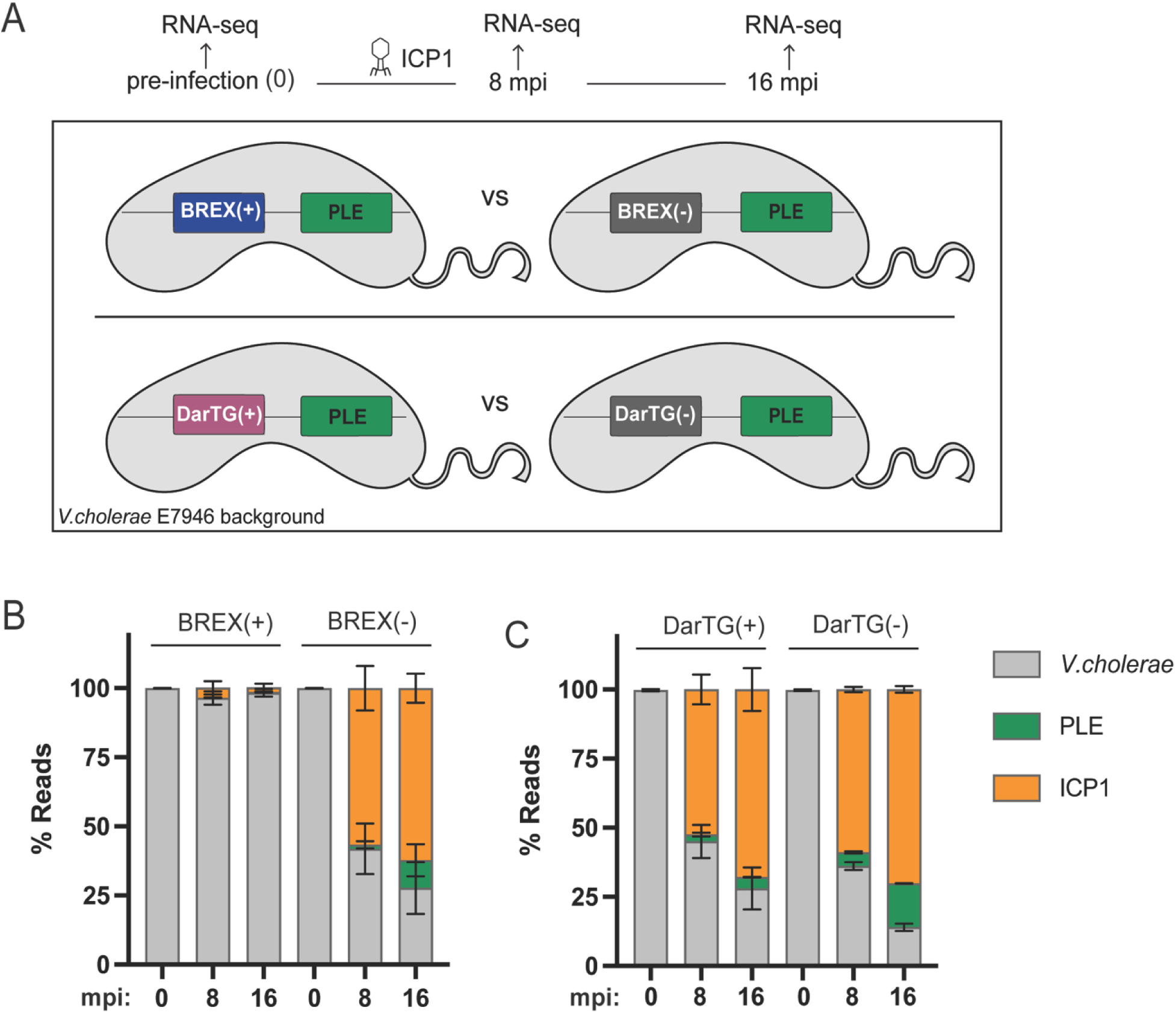
Experimental framework to evaluate the transcriptional impact of defense systems on phage and satellite gene expression. (A) Schematic of the RNA-seq experimental design. Samples were collected from *V. cholerae* E7946 PLE(+) strains before ICP1 infection (t=0), and at 8 and 16 minutes post-infection (mpi). Isogenic strain sets were compared to evaluate the impact of functional BREX and DarTG defenses (+) against their respective replication-permissive controls (-). (B) Proportional transcript abundance (percent reads) of the *V. cholerae* host, PLE, and ICP1 over the infection time course in the presence and absence of BREX. (C) Proportional transcript abundance (percent reads) of the host, PLE, and ICP1 over the infection time course in the presence and absence of DarTG. For both panels, the bar height represents the mean of three biological replicates; error bars indicate standard deviation.

To understand how these defense systems impact phage and satellite gene expression, we performed RNA sequencing (RNA-seq) on samples collected immediately before infection (0), during active viral replication in a productive infection (8 minutes), and late in the infection cycle (16 minutes) ^12^ (Fig. 1A). Principal component analysis confirmed distinct clustering by infection condition and genetic background (Fig. S1B). In the absence of BREX, we observed rapid transcriptional takeover by the phage and satellite: by 16 min post-infection (mpi), ICP1 and PLE transcripts constituted 62% and 11% of the total reads, respectively (Fig. 1B), closely mirroring previous analyses done in the same strain background without any additional defense-associated MGE ^12^. On the other hand, the presence of BREX severely limited this takeover, with host transcripts remaining dominant throughout the time course and ICP1 and PLE combined accounting for just 2% of total reads at 16 mpi (Fig. 1B). The absence of phage transcriptional takeover is consistent with the ability of BREX to block phage production while permitting cell survival ^3^. In stark contrast, DarTG did not repress the global accumulation of phage transcripts. At 16 mpi, ICP1 transcripts dominated the population equally in the presence or absence of the defense (67% and 70% of total reads, respectively; unpaired t-test *p* = 0.625) (Fig. 1C). However, while phage transcription was largely unaffected, PLE transcripts were significantly reduced in the presence of DarTG, accounting for only 4% of total reads compared to the 15% in the DarTG(-) control, (unpaired t-test *p* < 0.0001) (Fig. 1C).

### BREX potently arrests ICP1 gene expression, while DarTG permits replication-independent late gene expression

We next examined how these defenses alter the temporal progression of the phage life cycle. In the presence of BREX, ICP1 transcription was severely impaired. Visualization of coverage tracks revealed a global reduction in reads mapping to the phage genome compared to the permissive background (Fig. 2A, Fig. S2A). Differential expression analysis at 16 mpi showed that all phage transcripts were significantly downregulated (Fig. 2B, Tables S1 and S2). Analyses at both 8 and 16 mpi showed that this repression was most dramatic for the late-gene operons encoding capsid (*gp120*-*128*), and tail (*gp69-87* and *gp91-94*) morphogenesis proteins, as well as the holin/antiholin (*gp137-138*) lysis module (Fig. 2B). Interestingly, at 8 mpi, a small subset of immediate early genes (11 out of 227 genes) was not differentially regulated in the BREX(+) background. (Fig. 2B). This suggests that while BREX permits the initiation of the phage developmental program, it causes a premature arrest shortly after the onset of early transcription.

**Fig. 2:**
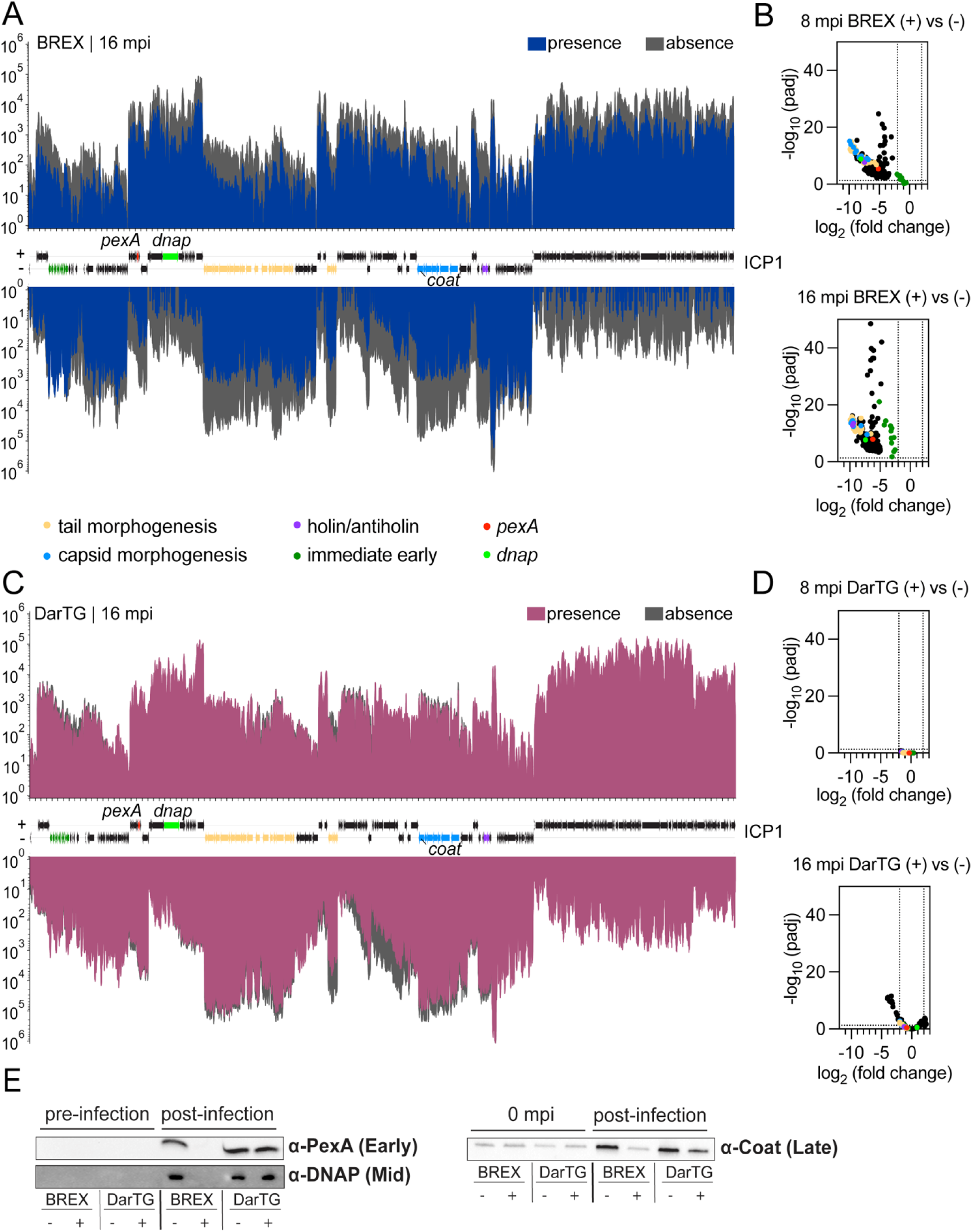
ICP1 gene expression progression is altered in the presence of anti-phage elements. (A) Average RNA-seq read coverage mapped to the ICP1 genome at 16 mpi in the presence or absence of BREX. Forward strand (+) reads are plotted above the x-axis and reverse strand (–) reads below the x-axis. Coverage represents the mean of three biological replicates and is displayed on a log-scaled y-axis. For A-D, ICP1 gene features are colored based on known or predicted functions according to the legend below panel A, while genes without functional annotation are shown in black. (B) Volcano plot illustrating the differential expression of ICP1 genes in the presence versus absence of BREX at 8 and 16 mpi. Vertical dotted lines indicate a log_2_ fold change cutoff of ≥ ±2, and horizontal dotted lines indicate a significance threshold of -log_10_ (p_adj_) ≥ 1.3. (C) Strand-specific RNA-seq coverage mapped to the ICP1 genome at 16 mpi in the presence or absence of DarTG. Plotting conventions are the same as in panel A. (D) Volcano plot illustrating the differential expression of ICP1 genes in the presence versus absence of DarTG at 8 and 16 mpi. Cutoff lines are the same as in panel B. (E) Western blots of ICP1 proteins probed in cells with and without BREX or DarTG. For the detection of PexA (early), DNAP (mid), and coat (late) proteins, samples were assessed at 8, 12, and 16 mpi, respectively. For the coat protein, the faint band appearing in infected BREX(+) cells originates from the initial phage inoculum. Samples at 0 mpi, (immediately upon addition of the phage inoculum) were probed to demonstrate that this faint band is a structural component of the input phage rather than *de novo* synthesis. The images are representative of three biological replicates. For the uncropped images of all replicates and quantification, see Fig. S3 and S4.

In stark contrast, visualization of coverage tracks revealed that DarTG had a minimal impact on the phage transcriptional program throughout the infection (Fig. 2C, Fig. S2B). At 8 mpi, no ICP1 genes were differentially regulated (Fig. 2D, Table S3). By 16 mpi, only a small subset of the phage genome (12 out of 227 genes, all of unknown function) was significantly downregulated (Fig. 2D, Table S4). The vast majority of the ICP1 transcriptome remained unaffected, indicating that, unlike BREX, DarTG allows the phage to execute a nearly complete temporal gene expression cascade despite blocking genome replication.

To determine if these transcriptional profiles were reflected at the protein level, we performed western blots for representative early (PexA), middle (DNA Polymerase, DNAP), and late (coat) phage proteins. Consistent with the severe repression observed by RNA-seq, no *de novo* synthesis of these proteins was detectable in the presence of BREX (Fig. 2E). This demonstrates that BREX, whose mechanism of defense remains unknown, restricts phage gene expression. Conversely, in the DarTG(+) background, all three phage proteins were readily detectable, confirming that the phage progresses to late-stage gene expression despite the replication block (Fig. 2E). However, densitometric quantification revealed nuances in the efficiency of this progression. While early and middle protein levels (PexA and DNAP, respectively) remained comparable to the replication-permissive control (Fig. S3A and S4A), we observed a significant approximately 2-fold reduction in the accumulation of the coat protein in the presence of DarTG (Fig. S3B). This protein-level decrease is consistent with a downward trend in *coat* transcription observed in our RNA-seq data (log_2_ fold change = -1.5 compared to DarTG(-)). However, this transcriptomic change did not meet the stringent significance thresholds required for global differential expression analysis.

Together, these results reveal that while both defense systems block phage genome replication, they impose vastly different constraints on phage gene expression. BREX potently arrests ICP1 gene expression shortly after entry. In contrast, DarTG allows the phage to produce late-stage structural proteins even in the absence of a replicated genome. This disparate effect also reveals a surprising uncoupling of late gene transcription from genome replication in ICP1, a finding that deviates from the canonical model for other phages ^24^ and some dsDNA eukaryotic viruses ^25^, in which viral genome replication is a prerequisite to license the activation of late gene expression.

### PLE gene expression is uncoupled from its genome replication

Given the divergent effects of BREX and DarTG on the helper phage’s gene expression, we hypothesized that these defenses would impact the PLE life cycle in distinct ways. We first examined PLE excision and circularization, the initial activation step triggered by the ICP1-encoded protein PexA^13^. We found that despite the presence of either BREX or DarTG defense systems, PLE successfully excises from the *V. cholerae* chromosome (Fig. 3A). While successful PLE excision was expected in the presence of DarTG, where PexA is expressed, it was surprising in the BREX(+) background, given that PexA was undetectable by western blot (Fig. 2E). This suggests that the threshold of PexA required to stimulate PLE excision is extremely low and falls below the limit of detection in our western blot analysis.

**Fig. 3:**
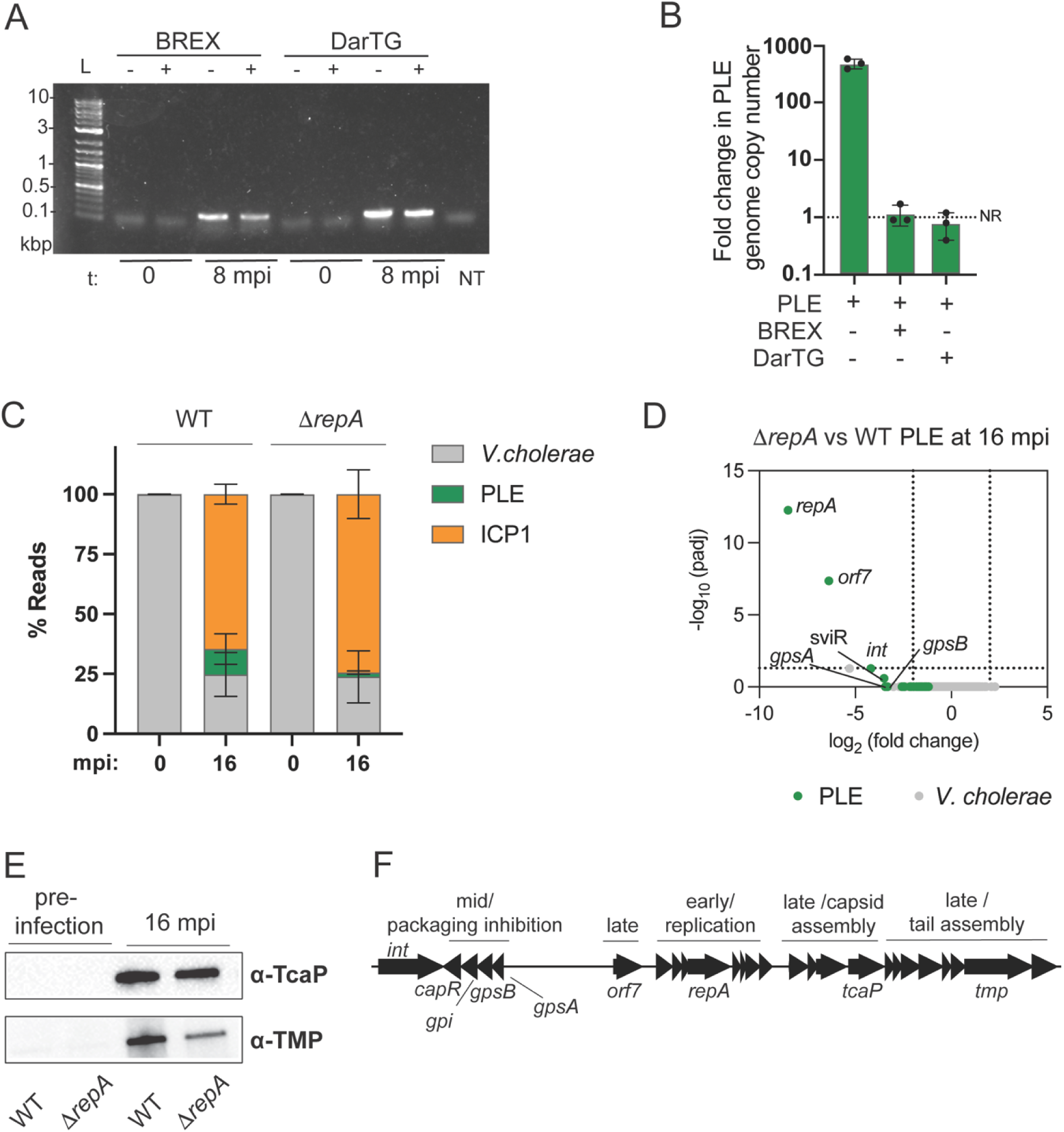
PLE transcriptional activation is uncoupled from genome replication. (A) PCR analysis of PLE circularization before (t=0) and 8 mpi in the presence (+) or absence (-) of the indicated defense system. L: Ladder, NT: No Template control. For additional biological replicates, see Fig. S5A. (B) Fold change in PLE genome copy number as assessed by qPCR at 20 mpi by ICP1. NR: No Replication. (C) Proportional transcript abundance (percent reads) of the *V. cholerae* host, PLE, and ICP1 before (0) and 16 mpi in WT PLE and the replication-deficient Δ*repA* mutant. (D) Volcano plot illustrating differential gene expression of Δ*repA* vs WT backgrounds at 16 mpi. Vertical dotted lines indicate a log_2_ fold change cutoff of ≥ ±2, and horizontal dotted lines indicate a significance threshold of -log_10_ (p_adj_) ≥ 1.3. (E) Representative western blots for PLE-encoded late proteins TcaP and TMP in WT and Δ*repA* backgrounds before (pre-infection) and 16 mpi by ICP1. For the uncropped images of all replicates and quantification, see Fig. S5B-D. (F) Schematic of the PLE1 genome. Genes are organized into temporal expression blocks (early, mid, late) based on peak expression during ICP1 infection ^12^, with corresponding functional modules indicated above.

Following excision, PLE replicates to high copy number, a process that remains incompletely understood, but is known to require the PLE-encoded replication initiation factor, RepA ^14^, and an ICP1-encoded helicase ^26^, and likely ICP1-encoded DNAP ^14^. With this in mind, we next measured the fold-change in PLE copy number by qPCR and found that PLE replication is completely blocked in the presence of BREX or DarTG (Fig. 3B). This observation prompted us to determine if PLE’s transcriptional activation is dependent on its own genome replication, as a block in this process could, in turn, be responsible for any observed effects on gene expression.

To directly test if PLE replication is a prerequisite for its transcriptional activation, we compared the transcriptional profiles of wild-type (WT) PLE and the replication-deficient mutant (Δ*repA*) at 16 mpi by RNA-seq. Strikingly, principal component analysis showed that WT and Δ*repA* conditions cluster similarly during infection (Figure S1B). In the WT background, PLE reads accounted for 10.6% of the total transcriptome, whereas in the Δ*repA* mutant, PLE transcripts accounted for 1.7% (unpaired t-test *p* = 0.134), a comparably high level of expression compared to the 0.03% observed in uninfected cells (Fig 3C). Differential gene expression analysis confirmed only *orf7* was significantly downregulated in the Δ*repA* mutant (Fig. 3D). While a subset of the remainder of the PLE genome (Fig. 3F)—including the integrase (*int*), the packaging inhibition operon (*gpsAB, gpi,* and *capR*) ^16^, and the small non-coding RNA SviR ^27^—exhibited a trend toward lower expression in the Δ*repA* mutant (with log_2_ fold change < - 3), they did not meet the stringent significance thresholds required to be defined as significantly downregulated (Fig. 3D and Table S5).

To validate this observed transcriptional activation of PLE at the protein level, we assessed the production of TcaP ^15^ and the tape measure protein (TMP) ^27^, two late-stage ^12^ PLE-encoded structural proteins involved in the formation of PLE transducing particles. Consistent with the RNA-seq data, both proteins were produced even in the absence of PLE replication (Fig. 3E). However, while TcaP levels were comparable to WT, TMP abundance was significantly reduced in the Δ*repA* mutant (Fig. S5B-D). Since *tmp* transcripts were not significantly downregulated (Table S5), this reduction likely reflects post-transcriptional regulation, potentially linked to the lower abundance of SviR. SviR is known to increase the transcript stability of the *capR* operon ^28^; correspondingly, this operon also trended toward lower expression in the Δ*repA* mutant (log_2_ fold change -3.3). Altogether, these data demonstrate that PLE’s robust transcriptional activation during ICP1 infection—including the production of late-stage structural proteins—occurs independently of its genome replication. This aligns with our previous findings that PLE replication is not required for the functional inhibition of ICP1^14,26^.

### BREX stalls PLE gene expression

We reasoned that because the anti-phage defense systems BREX and DarTG disrupt the progression of ICP1 gene expression to different extents, they could serve as tools to probe how PLE’s transcriptional program responds to perturbations in the ICP1 life cycle. Since we established that PLE replication is not necessary for its transcriptional activation (Fig. 3), we investigated the transcriptional activity of PLE in the presence and absence of BREX in greater detail.

First, we examined the read coverage plots of PLE in the BREX(-) background. In the absence of BREX, read coverage plots demonstrate that, as expected, the entire PLE transcriptional program was robustly induced upon ICP1 infection (Fig. 4A). Differential expression analysis revealed that all PLE genes were upregulated at 8 and 16 mpi compared to uninfected cells, with expression levels increasing as the infection progressed (Fig. 4C, Fig. S6A, and Tables S6 and S7).

**Fig. 4:**
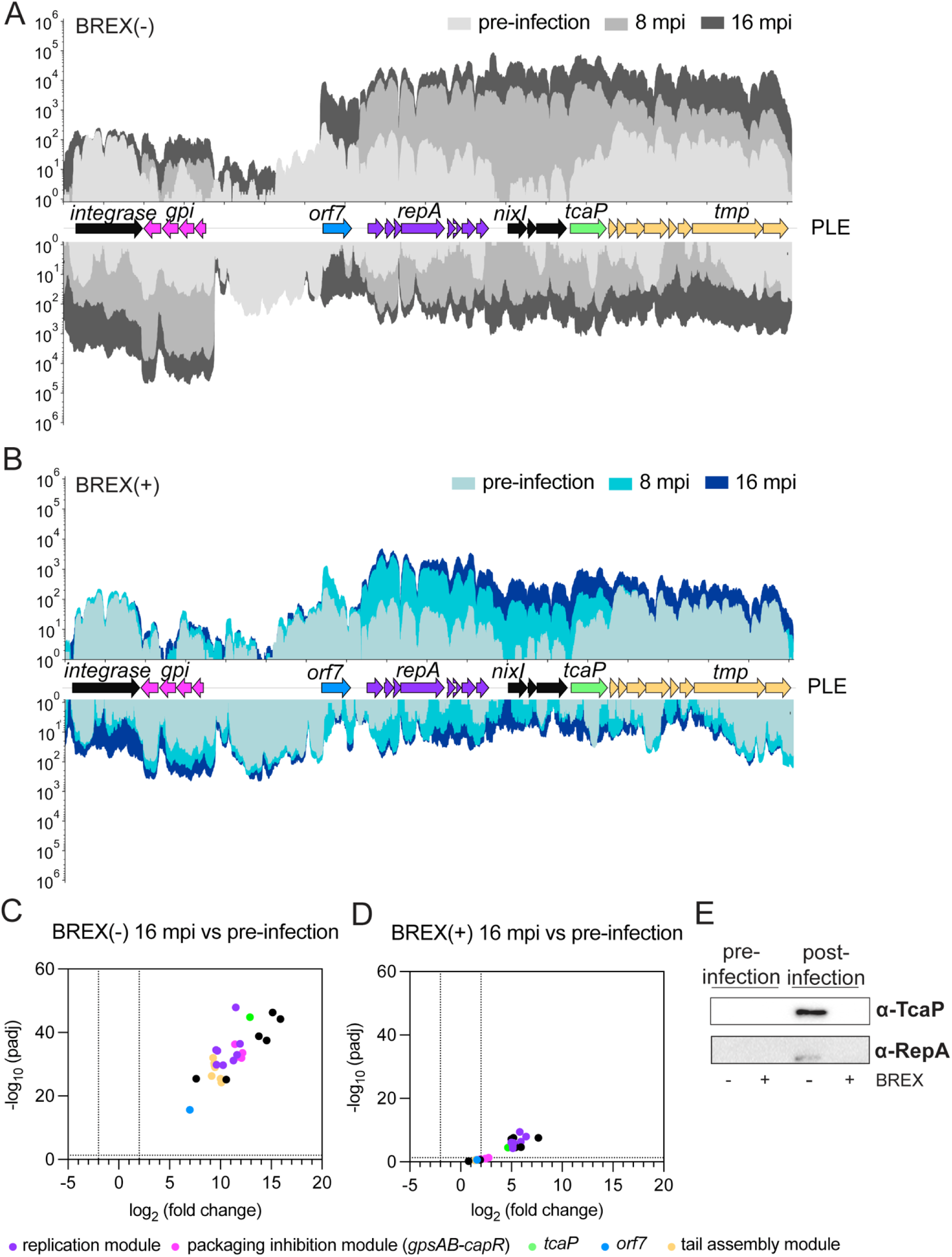
BREX stalls the PLE transcriptional program. (A, B) Average RNA-seq read coverage mapped to the PLE genome in (A) BREX(-) and (B) BREX(+) cells. Coverage tracks represent uninfected, 8 mpi, and 16 mpi timepoints. Forward-strand reads are plotted above the x-axis and reverse-strand reads below. Coverage represents the mean of three biological replicates (with the exception of two biological replicates for BREX(+) pre-infection sample, see methods for detail) and is displayed on a log-scaled y-axis. (C, D) Volcano plots illustrating differential PLE gene expression at 16 mpi compared to pre-infection in (C) BREX(-) and (D) BREX(+) backgrounds. Each point represents a PLE gene. Selected genes belonging to the functional modules defined in Fig. 3F are colored according to the legend, while other PLE genes are shown in black. Vertical dotted lines indicate a log_2_ fold change cutoff of ≥ ± 2, and horizontal dotted lines indicate a significance threshold of -log_10_ (p_adj_) ≥ 1.3. (E) Western blots for PLE-encoded TcaP (late) and RepA (early) probed in cells with (+) and without (-) BREX before and after infection. TcaP was probed at 16 mpi and RepA at 12 mpi. Images are representative of three biological replicates. For the uncropped images of all replicates, see Fig. S7.

In stark contrast, read coverage plots of PLE in the BREX(+) background showed that gene expression remained minimal throughout the course of infection (Fig. 4B). However, differential expression analysis comparing infected to uninfected BREX(+) cells revealed that a subset of genes, comprised of the early replication operon (*orf8* to *orf14)* and *nixI-tcaP*, is upregulated during infection (Fig. 4D, Fig. S6A, and Tables S8 and S9). While this specific subset of genes is upregulated, the magnitude of induction is drastically lower than in the BREX(-) background. For example, at 16 mpi, *repA* and *tcap* showed only 35- and 25-fold increases in BREX(+) cells, respectively, compared to >1,000- and >7,000-fold increases in the BREX(-) background. Furthermore, the remaining PLE modules, including the packaging inhibition operon (*gpsAB* and *gpi*), *capR, orf7,* and the tail assembly operon (*orf18-orf23*), did not show any significant upregulation in the presence of BREX. This suggests that PLE activation requires multiple distinct cues. In the BREX(+) background, at least some of these cues are likely disrupted or limited, preventing complete and robust transcriptional activation of the satellite.

To validate these findings at the protein level, we assessed the production of the PLE-encoded early protein RepA and the late protein TcaP. Strikingly, despite the significant >20-fold increase in transcript levels observed for both genes, neither protein was detectable by western blot in the presence of BREX (Fig. 4E). This reveals a notable disconnect between transcript accumulation and protein production. While the absolute abundance of these transcripts in the BREX(+) background may remain below a required threshold for robust translation, particularly when compared to the orders-of-magnitude higher induction seen in the BREX(-) background, this result strongly suggests that BREX may impose an additional block on gene expression post-transcriptionally. Alternatively, the premature arrest of the ICP1 life cycle by BREX may deprive PLE of specific helper-encoded factors required for efficient translation or stability of its own proteins. Regardless of the mechanism, the absence of RepA directly aligns with the complete block in PLE replication observed in these cells (Fig. 3B). Overall, these results demonstrate that while PLE can initiate the first steps of its transcriptional program, BREX effectively stalls the satellite life cycle before proteins can be produced.

### DarTG permits robust PLE gene expression

In contrast to BREX, DarTG had a much milder effect on the overall transcriptional landscape of PLE (Fig. 5A and 5B). PLE is robustly activated in the presence of DarTG, despite the block in replication (Fig. 5B, 3B). Read coverage and differential expression analyses showed that the entire PLE transcriptional program was induced, although the magnitude of upregulation was slightly lower than in the DarTG(-) control (Fig. 5C and 5D). This observation is corroborated by the global ∼4-fold reduction in PLE transcripts, from 15% to 4% of the total transcriptome (Fig. 1C), which may reflect PLE’s replication deficiency in the presence of DarTG (Fig. 3B).

**Fig. 5:**
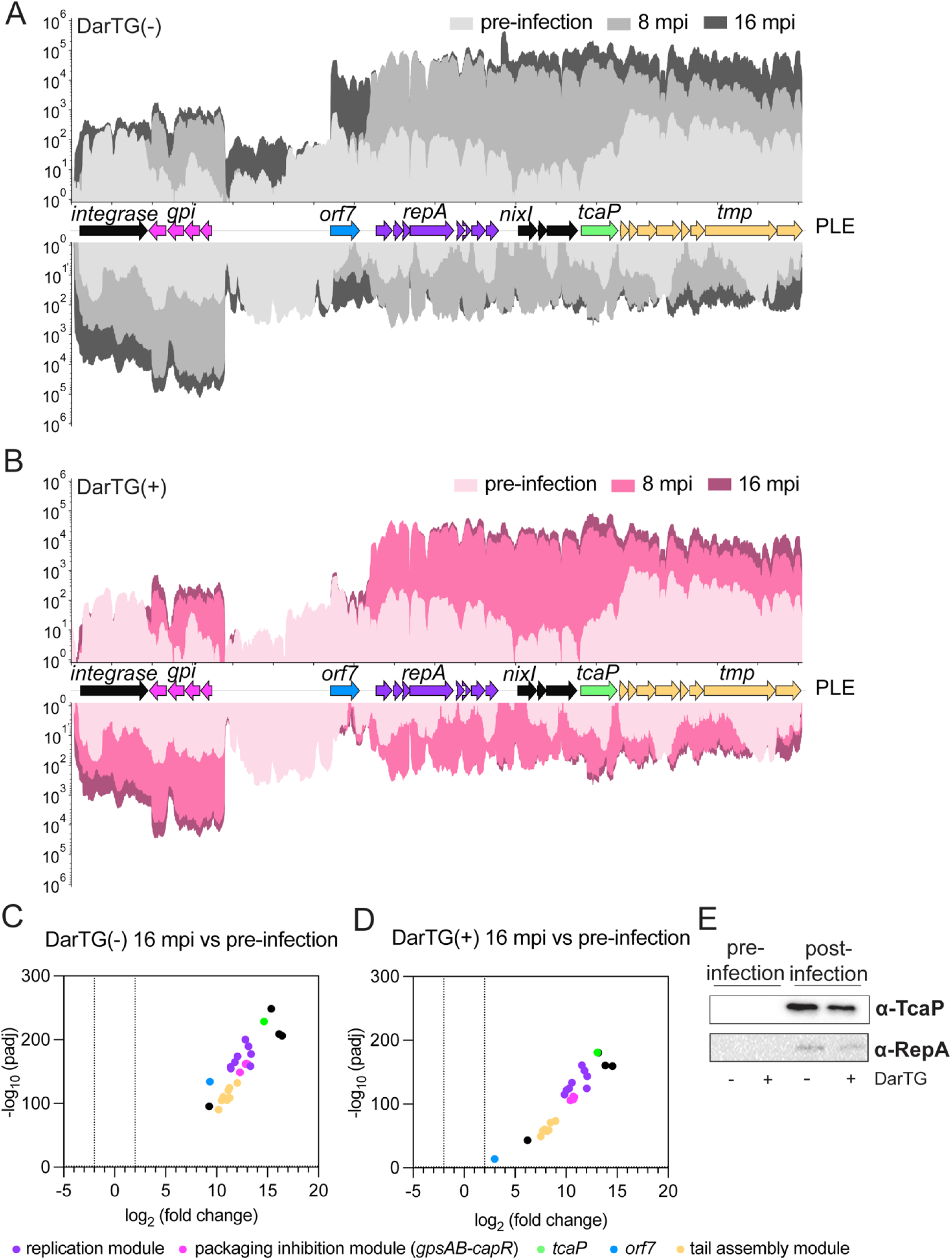
DarTG permits the progression of the PLE transcriptional program. (A, B) Average RNA-seq read coverage mapped to the PLE genome in (A) DarTG(-) and (B) DarTG(+) cells. Coverage tracks represent uninfected, 8 mpi, and 16 mpi timepoints. Forward-strand reads are plotted above the x-axis and reverse-strand reads below. Coverage represents the mean of three biological replicates and is displayed on a log-scaled y-axis. (C, D) Volcano plots illustrating differential PLE gene expression at 16 mpi compared to pre-infection in (C) DarTG(-) and (D) DarTG(+) cells. Each point represents a PLE gene. Selected genes belonging to the functional modules defined in Fig. 3F are colored according to the legend, while other PLE genes are shown in black. Vertical dotted lines indicate a log_2_ fold change cutoff of ≥ ± 2, and horizontal dotted lines indicate a significance threshold of -log_10_ (p_adj_) ≥ 1.3. (E) Western blots for PLE-encoded TcaP (late) and RepA (early) probed in cells with (+) and without (-) DarTG before and after infection. TcaP was probed at 16 mpi and RepA at 12 mpi. Images are representative of three biological replicates. For the uncropped images of all replicates, see Fig. S7.

At 8 mpi, all PLE genes appeared to be upregulated to a similar magnitude in both DarTG(+) and DarTG(-) cells compared to uninfected controls (Fig. S6B and Tables S10, S12). However, by 16 mpi, the tail morphogenesis operon (*orf18*-*orf23*) was more potently upregulated in the absence of DarTG (Fig. 5C and 5D, Tables S11 and S13). For example, *tmp* showed a ≈ 230-fold increase in the presence of DarTG, compared to a ≈ 2,300-fold increase in its absence. Notably, *orf7* showed the highest magnitude difference in induction, with an 8-fold increase in DarTG(+) cells compared to the >640-fold increase in the control. This specific difference can be attributed to the PLE’s inability to replicate in the presence of DarTG, as *orf7* was the only gene significantly downregulated in the Δ*repA* mutant (Fig. 3D). In contrast, at 16 mpi, PLE genes within the replication module (*orf8*-*orf13*) and the packaging inhibition module (*gpsAB*-*capR*) reached similar magnitudes of induction in both backgrounds. For example, *repA* increased ≈1,500-fold in the presence of DarTG and ≈4,200-fold in its absence (Fig. 5C and 5D).

Consistent with these transcriptional profiles, PLE gene expression culminated in the production of both the early protein RepA and the late protein TcaP (Fig. 5E). These results indicate that, similar to our observations for ICP1, the production of PLE-encoded late structural proteins does not strictly depend on PLE genome replication (Fig. 5E). Although RepA protein was detectable, PLE replication remained blocked (Fig. 3B). This is consistent with the known activity of DarT, which ADP-ribosylates single-stranded DNA to inhibit DNA synthesis ^29,30^. Together, these results show that, unlike the early arrest of the ICP1 life cycle in the presence of BREX, DarTG allows the ICP1 life cycle to progress sufficiently to provide the necessary cues for the progressive activation of the PLE transcriptional program.

## DISCUSSION

In this study, we used BREX and DarTG anti-phage defense systems as tools to probe the regulatory logic of PLE gene expression. We initially established that, while both systems completely block the ICP1 helper phage’s genome replication, they exert distinct effects on the phage’s transcriptional program. BREX imposes a profound arrest, restricting ICP1 gene expression to a small subset of immediate-early genes and preventing the accumulation of mid and late-stage proteins. In contrast, DarTG is relatively permissive, allowing the phage transcriptional program to proceed almost entirely unperturbed. This culminates in the production of early, mid, and late proteins despite the block in DNA synthesis. The latter finding was somewhat surprising, given previous data showing that plasmid-based expression of DarTG in *Escherichia coli* substantially reduced RNA synthesis following phage infection ^31^. Because the system evaluated here is in its native genetic context, our experiments may have better captured physiologically relevant gene expression dynamics, although the specific molecular roadblocks imposed by host defenses may inherently vary across different phage-host pairs.

The contrasting transcriptional landscapes imposed by these defense systems provided a unique opportunity to investigate PLE regulation, as neither background permits PLE replication. To untangle whether the resulting PLE transcriptional profiles were a consequence of this shared replication deficiency, we characterized the transcriptome of a replication-deficient (Δ*repA*) PLE mutant. Our results revealed that PLE still robustly expresses nearly its entire transcriptional program even without replicating, demonstrating that PLE gene expression and structural protein production are effectively uncoupled from its own replication. Hence, upon excision from the host chromosome to escape chromosomal degradation ^26^, a single episomal copy of PLE is sufficient to execute a robust gene expression program. Curiously, *orf7*, encoding a predicted MarR-like transcription factor, emerged as the only PLE gene significantly impacted by PLE’s replication status. Although the function of ORF7 remains unknown and it is dispensable for ICP1 inhibition ^32^, its unique dependence on replication suggests it may play a regulatory role in the late stages of infection, perhaps optimizing PLE horizontal transfer.

In the presence of BREX, we observed a transcriptional stall in PLE gene expression that mirrors the arrested state of the ICP1 helper phage developmental program. Previous studies suggest that BREX acts early during infection, primarily by blocking phage DNA replication ^3^, though the precise mechanism of phage restriction remains unclear. To our knowledge, no prior studies have investigated whether BREX impacts gene expression of the invading phage. We observed that in the presence of BREX, ICP1 initiates expression of immediate-early genes but fails to reach full early-, mid-, or late-stage expression. This early window of infection appears to trigger modest transcriptomic induction of the PLE replication module; however, most of the remaining PLE modules remain transcriptionally silent in the BREX(+) background, suggesting that the cues required to fully license robust transcription are produced only as the phage lifecycle advances.

A central question is how PLE senses the progression of its helper phage to initiate transcription. While PLE may directly respond to phage-encoded proteins, our data are also consistent with a model in which activation depends on modulation of host factors during progression through the ICP1 lifecycle. The global host regulator H-NS is the primary repressor of PLE gene expression ^21^. PLE transcriptional activation likely involves antagonizing this repression, though our findings indicate that this derepression is not a singular, all-or-none event. Instead, the sequential activation of PLE operons suggests that each operon is derepressed as distinct milestones of the ICP1 developmental program are achieved.

The progressive licensing of PLE during ICP1 infection represents a substantial departure from the canonical regulatory architecture exemplified by SaPIs. In the SaPI model, a single non-essential protein typically suffices to relieve SaPI repression and trigger the entire lifecycle ^19,20^. Although, to our knowledge, global assessments of satellite activation during helper phage infection have not been carried out outside the PLE-ICP1 system, there is evidence from another satellite that reliance on the helper can be more modular and distributed than the SaPI one anti-repressor/one repressor paradigm. Specifically, activation of the unrelated satellite P4 requires two distinct helper-encoded inputs for full induction by its cognate phage ^33,34^. Independent gating of replication and morphogenesis operons in satellites may ensure that replication functions are rapidly available upon infection, whereas structural components are expressed only once the helper phage has progressed to a stage compatible with satellite packaging and transfer. Such a multi-layered architecture is likely less vulnerable to phage escape through simple bypass of a single sensing mechanism.

## MATERIALS AND METHODS

### Generation of appropriate controls for the experimental design

Two anti-phage defense systems that block the initiation of ICP1 genome replication in clinical isolates of *V. cholerae* were used in this study: BREX, encoded by *Vch*Ind5, an integrative and conjugative element ^22^, and DarTG, a toxin-antitoxin system encoded by a distinct phage defense element ^23^. To enable appropriate comparisons, we separately introduced the genetic elements encoding BREX and DarTG into the same parental *V. cholerae* strain harboring PLE1 (see conjugations and transductions, below). To generate isogenic controls, we deleted hotspot five (encoding BREX) from *Vch*Ind5 ^22^, and designated it as BREX(-). As a permissive background for phage defense element encoding DarTG, we deleted *darT* (the toxin, shown to inhibit ICP1 genome replication in its native genetic context ^23^) and also *old*, encoded by the same element but only shown to inhibit ICP1 when expressed from a plasmid ^23,35^), and designated the resulting strain as DarTG(-). For all infections, we used a Δ*orbA* derivative of ICP1_2006_Dha_E ΔCRISPR-cas Δ*cas2-3*, as OrbA is the known inhibitor of the *Vch*Ind5 BREX system ^22^. The deletion of the ICP1 CRISPR-Cas system was necessary to prevent ICP1-mediated anti-PLE activity.

### Bacterial strains and growth conditions

Bacterial strains used in this study are listed in Table S14. All *V. cholerae* and *E. coli* strains were grown with aeration in a roller at 250 rpm at 37°C in Lysogeny Broth (LB). Where necessary, antibiotics were used at the following concentrations: 100 μg/mL streptomycin, 75 μg/mL kanamycin, 100 μg/mL spectinomycin, 32 μg/mL trimethoprim, 1.25 μg/mL chloramphenicol (*V. cholerae* in liquid), 2.5 μg/mL chloramphenicol (*V. cholerae* on plates), 25 μg/mL chloramphenicol (*E. coli*). For induction of plasmids in *V. cholerae*, 1 mM isopropyl-β-D-thiogalactopyranoside (IPTG) and 1.5 mM theophylline were added when cultures reached at an optical density (OD_600_) of 0.2 and were grown for 20 additional minutes at 37°C in the roller. For induction of *tcaP* expression in *E. coli* as a positive control for western blotting, 1mM IPTG was added to cultures at an OD_600_ of 0.2 and cultures were grown for an additional 20 minutes at 37°C in the roller.

### Generation of mutant strains

Chromosomal deletions in *V. cholerae* were constructed by PCR amplifying 1kb upstream and downstream homology arms, which then joined with a spectinomycin resistance cassette flanked by FRT (flippase recognition target) sites using splicing by overlap extension PCR. The stitched PCR product containing the arms of homology and the selective marker was then introduced into *V. cholerae* by natural transformation. To achieve natural competency, cells were grown in LB at 30°C until OD_600_=0.3. The cells were then pelleted and washed twice with 0.5x Instant Ocean. Then, cells were added to 80 mg chitin (Sigma) in 0.5x Instant Ocean and incubated for 24 hours at 30°C. The transformants were plated on LB agar containing antibiotic. To remove the spectinomycin cassette, the inducible plasmid encoding the flippase was conjugated from *E. coli* S17 into *V. cholerae* and exconjugants were selected on streptomycin and chloramphenicol. Flippase expression was induced using 1 mM IPTG with 1.5 mM theophylline. Removal of the spectinomycin cassette and loss of the plasmid were verified by plating on LB agar plates supplemented with spectinomycin and chloramphenicol. Colonies that did not grow on spectinomycin or chloramphenicol plates were purified on LB agar supplemented with streptomycin and the deletion was confirmed by PCR and Sanger sequencing.

### Conjugations and transductions

To obtain PLE(+) *Vch*Ind5(+) cells, *V. cholerae* E7946 PLE(+) was conjugated with *V. cholerae Vch*Ind5(+) donors (wild-type and the Δhotspot 5 BREX(-) derivative). *Vch*Ind5 donor *V. cholerae* strains were grown overnight with aeration in a roller at 250 rpm at 37°C in LB medium supplemented with trimethoprim, and PLE (+) recipient strains were grown with kanamycin. 500 μL of overnight-grown donor and recipient cultures were mixed and pelleted at 5,000 × *g* for three minutes. Pellets were resuspended in 50 μL LB and pipetted onto an 0.8 μm filter disk on an LB agar plate. Conjugations were allowed to proceed for 6 hours at 37°C. After the incubation, the filter disk was removed and added to a microcentrifuge tube with 1 mL LB. The microcentrifuge tube was vigorously vortexed to allow bacteria to dislodge from the filter into the liquid LB, and a dilution series was plated on trimethoprim and kanamycin to select for PLE(+) recipients that acquired *Vch*Ind5. Exconjugants were then purified further on trimethoprim and kanamycin plates two more times.

To obtain the PLE(+) DarTG(+) strain, PLE was transduced into the recipient DarTG(+) *V. cholerae* ^23^. Donor PLE(+) *V. cholerae* was grown with aeration in a roller at 250 rpm at 37 °C in LB to OD_600_=0.3. Recipient DarTG(+) cells were grown overnight in the same conditions. Donor PLE(+) cells were infected with ICP1 at an MOI of 2.5, then the culture was washed twice to remove unbound phage and resuspended in LB with 10 mM MgCl_2_. After lysis, the culture was treated with chloroform and the resulting lysate was centrifuged at 5,000 × *g* for 15 minutes at 4 °C. The supernatant, which contains the PLE transducing particles, was added to the overnight recipient culture supplemented with 10 mM MgCl_2_ in a 1:1 ratio and incubated for 20 minutes with aeration at 250 rpm at 37 °C. Transductants were plated on kanamycin and spectinomycin to select for recipients that acquired PLE. To obtain the otherwise isogenic PLE(+) DarTG(-) strain, *darT* and *old* were deleted in this background as described above.

### Quantification of genome replication by quantitative polymerase chain reaction (qPCR)

We used an established protocol to quantify phage genome replication as previously described ^11^. Briefly, *V. cholerae* strains were grown on a plate with an appropriate antibiotic overnight at 37 °C. The next day, the strain was grown to an OD_600_=1 in liquid culture. Then, the liquid culture was back-diluted to 0.05 and grown to 0.3 and infected with ICP1 Δ*orbA* strain at a multiplicity of infection (MOI) of 0.1. Immediately after infection (t=0), 100 μL samples were collected and boiled at 95°C for 10 minutes. The remaining culture was incubated at 37°C with aeration for 20 minutes (t=20), 100 μL of the culture was collected, and boiled. Phage DNA from boiled samples was diluted 20-fold and measured by qPCR with primers zac68 (CTGAATCGCCCTACCCGTAC) and zac69 (GTGAACCAACCTTTGTCGCC) using iQ SYBR Green Supermix (Bio-Rad) and the CFX Connect Real-Time PCR Detection system (Bio-Rad). DNA replication was determined by quantifying the fold change in replication at t=20 compared to genome copy at t=0 using the *C_q_* value relative to that of a standard curve of known concentrations of ICP1 genomic DNA. PLE replication followed the same protocol as above, with the following exceptions: t=0 is collected immediately before infection, strains were infected with ICP1 at an MOI=2.5, samples were diluted 500-fold before being used as templates for qPCR reactions, and primers zac14 (AGGGTTTGAGTGCGATTACG) and zac15 (TGAGGTTTTACCACCTTTTGC) specific for the PLE genome were used. For both phage and PLE genome replication, all conditions were tested in biological triplicate, and each reported data point is the mean of three technical replicates.

### Western blot analysis

*V. cholerae* strains were grown on a plate with an appropriate antibiotic overnight at 37 °C. The next day, the strain was grown to an OD_600_=1 in liquid culture. Next, the liquid culture was back-diluted to an OD_600_=0.05 and grown to an OD_600_=0.3. Then, a 1 mL aliquot of the culture was spun down at 5,000 × *g* for 3 minutes. The remaining culture was infected with ICP1 at an MOI of 2.5. For each time point post-infection indicated in the figure, a 1 mL aliquot of culture was collected and mixed with ice-cold methanol and centrifuged at 5,000 × *g* for 3 minutes. Cell pellets were resuspended in Laemmli buffer supplemented with β-mercaptoethanol (Bio-Rad). Samples were boiled for 10 minutes at 99 °C before loading 10 μL onto the Mini-Protean TGX Stain Free Precast PAGE gels (Bio-Rad). For quantifying the relative abundance of the protein of interest in different conditions, total protein was quantified by activating stain-free gels for 2 minutes on the UV/Stain-Free tray and imaged on Gel Doc EX Imager (Bio-Rad) and protein band intensities were measured using Fiji software (Version 2.16.0). The gel was then transferred to a mini-size PVDF membrane (Bio-Rad) via the Trans-Blot Turbo system. Proteins were detected via custom primary peptide antibodies generated in rabbits against PLE encoded TcaP, RepA, or ICP1 encoded DNAP, PexA, and Gp122 (GenScript). Primary antibodies were used at the following dilutions: RepA and Gp122 (1:1,500), PexA (1:15,000), DNAP (1:1,000), and TcaP (1:600). Secondary detection was performed using goat anti-rabbit IgG horseradish peroxidase antibody (Bio-Rad, diluted 1:10,000). The membrane was then developed with Clarity Western ECL substrate (Bio-Rad) and imaged on a ChemiDoc XRS imaging system (Bio-Rad). The intensity of the band corresponding to the protein of interest (verified by size and appropriate controls as indicated on the blots) was normalized to the total protein loaded to each well.

### RNA isolation for RNA-seq

Three biological replicates of RNA isolations were conducted for each background. *V. cholerae* strains were grown on a plate with an appropriate antibiotic overnight at 37 °C. The next day, the strain was grown to an OD_600_=1 in liquid culture. Next, the liquid culture was back-diluted to an OD_600_=0.05 and grown to an OD_600_=0.3. A four mL sample was removed before infecting with ICP1 at an MOI of 2.5, and four mL samples were collected at 8 and 16 mpi; all samples were immediately mixed with an equal volume of ice-cold methanol and pelleted at 5,000 × *g* for 5 minutes at 4 °C. The supernatant was removed, and the pellets were washed with ice-cold 1x phosphate-buffered saline (pH 6.2). The mixture was pelleted at 5,000 × *g* for 5 minutes at 4 °C, the supernatant was removed, and pellets were resuspended in 200 μL of TRIzol Reagent (Invitrogen). The samples were stored at -80 °C. To extract the RNA, the samples were thawed at room temperature (RT) for five minutes and 40 μl chloroform was added to each sample.

Samples were vigorously vortexed and incubated at RT for another 10 minutes. Then, to separate the phases, samples were centrifuged at 12,000 × g for 10 minutes at 4 °C. The top aqueous layer was mixed thoroughly with 110 μL of isopropanol and 11 μL of 3 M sodium acetate (pH 7.4) and incubated at RT for 10 minutes. Samples were then centrifuged at 12,000 × *g* for 15 minutes at 4 °C. The supernatant was discarded, and the pellet containing the RNA was washed twice with 500 μL 75% ethanol. To evaporate the remaining ethanol, the pellets were incubated at 65 °C for at least two minutes. Finally, RNA pellets were resuspended in 20 μL of RNAse-free water.

### RNA-seq analysis

Isolated RNA samples were sequenced by the SeqCenter (Pittsburgh, PA) with a sequencing depth of 12 million reads. The paired-end FASTQ files were mapped to the reference genomes using bowtie2 ^36^. Reference genomes were generated *in silico* as required to generate the appropriate *V. cholerae* chromosomes 1 (CP024162.1) and 2 (CP024163.1) and introducing the integrative conjugative element SXT *Vch*Ind5 (GQ463142) or phage defense element encoding DarTG ^23^. The resulting mapped alignment files were converted to BAM format, sorted, and indexed using samtools ^37^. Transcript abundance was estimated using StringTie ^38^. Transcript count matrices were generated using the prepDE.py script provided with StringTie. The matrices were imported into RStudio for downstream differential expression analysis using DESeq2 ^39^. Variance-stabilizing transformed expression values were used for PCA to assess data quality and sample clustering. One biological replicate for BREX(+) cells in the uninfected condition clustered aberrantly in the PCA plot and was excluded as an outlier for the comparison between BREX(+) infected and BREX(+) uninfected samples. Therefore, differential expression analysis was performed using two biological replicates for the BREX(+) uninfected condition, while three biological replicates were used for all other conditions and comparisons. Results were illustrated using volcano plots generated using GraphPad (version 10.5.0 (673)).

Sorted and indexed BAM files were used to generate strand-specific RNA-seq coverage tracks. Normalization factors were calculated using multiBamSummary, and strand-specific coverage tracks were generated using bamCoverage from deepTools ^40^. Coverage tracks were exported as BedGraph files and merged across biological replicates using unionbedg from BEDTools ^41^ to compute mean coverage for each condition. The resulting normalized strand-specific coverage tracks were visualized across ICP1 and PLE genomes together with gene annotations from the GTF file.

The proportion of transcript abundance for each element in the sample (percent reads) was calculated by normalizing raw integer counts to the total library size. Reads mapping to PLE, ICP1, and the host genome were aggregated by element and divided by the total library reads and expressed as a percentage of the total reads per sample. Results were illustrated using comparative bar graphs using GraphPad (version 10.5.0 (673)).

## Data availability

Sequence data for samples used in this work will be released upon publication and can be found in the Sequence Read Archive under the BioProject accession PRJNA1439424.

## Acknowledgements

We thank all Seed Lab members, past and present, for useful discussions, with special thanks to Drew Dunham for his support in the initial implementation and troubleshooting of our bioinformatics workflow. The project described was supported by Grant number R01AI127652 to K.D.S from the National Institute of Allergy and Infectious Diseases of the National Institutes of Health. Its contents are solely the responsibility of the authors and do not necessarily represent the official views of the National Institutes of Health. This work was supported in part by funding to K.D.S. as a Biohub, San Francisco, Investigator and by the Miller Institute for Basic Research in Science, University of California Berkeley. K.D.S. holds an Investigators in the Pathogenesis of Infectious Disease Award from the Burroughs Wellcome Fund.

