## Supplementary material for "Repurposing anti-phage defenses to differentially arrest the viral lifecycle reveals the regulatory logic of a parasitic satellite": Fig S1-S7 and Table S14

A

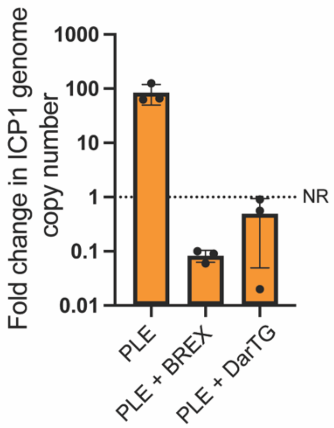
Fold change in ICP1 genome copy number
1000
100
10
1
0.1
0.01
NR
PLE
PLE + BREX
PLE + DarTG

B

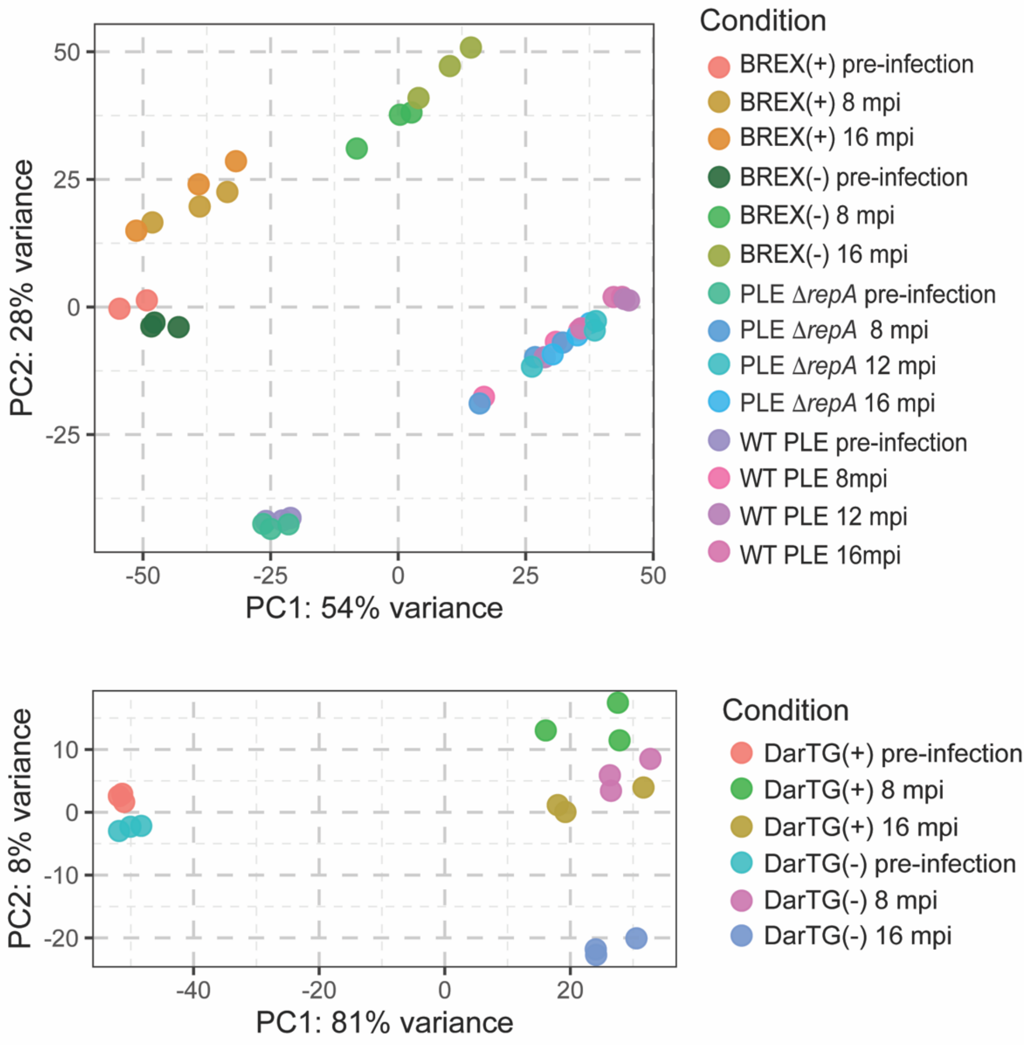
PC2: 28% variance
PC1: 54% variance
50
25
0
-25
-50
Condition
BREX(+) pre-infection
BREX(+) 8 mpi
BREX(+) 16 mpi
BREX(-) pre-infection
BREX(-) 8 mpi
BREX(-) 16 mpi
PLE ΔrepA pre-infection
PLE ΔrepA 8 mpi
PLE ΔrepA 12 mpi
PLE ΔrepA 16 mpi
WT PLE pre-infection
WT PLE 8mpi
WT PLE 12 mpi
WT PLE 16mpi
PC2: 8% variance
PC1: 81% variance
10
0
-10
-20
-40
-20
0
20
Condition
DarTG(+) pre-infection
DarTG(+) 8 mpi
DarTG(+) 16 mpi
DarTG(-) pre-infection
DarTG(-) 8 mpi
DarTG(-) 16 mpi

Fig. S1: (A) ICP1 genome replication assessed by qPCR at 20 mpi relative to 0 mpi in cells with PLE only, PLE and BREX, or PLE and DarTG. NR: No Replication (B) PCA was performed to reduce the dimensionality of the transcript count matrix in each condition. Each dot represents a biological replicate, and conditions are color-coded (N=3 for each condition, with the exception of the BREX(+) pre-infection, which was N=2 as one replicate was detected as an outlier during quality control and excluded from the downstream analysis).

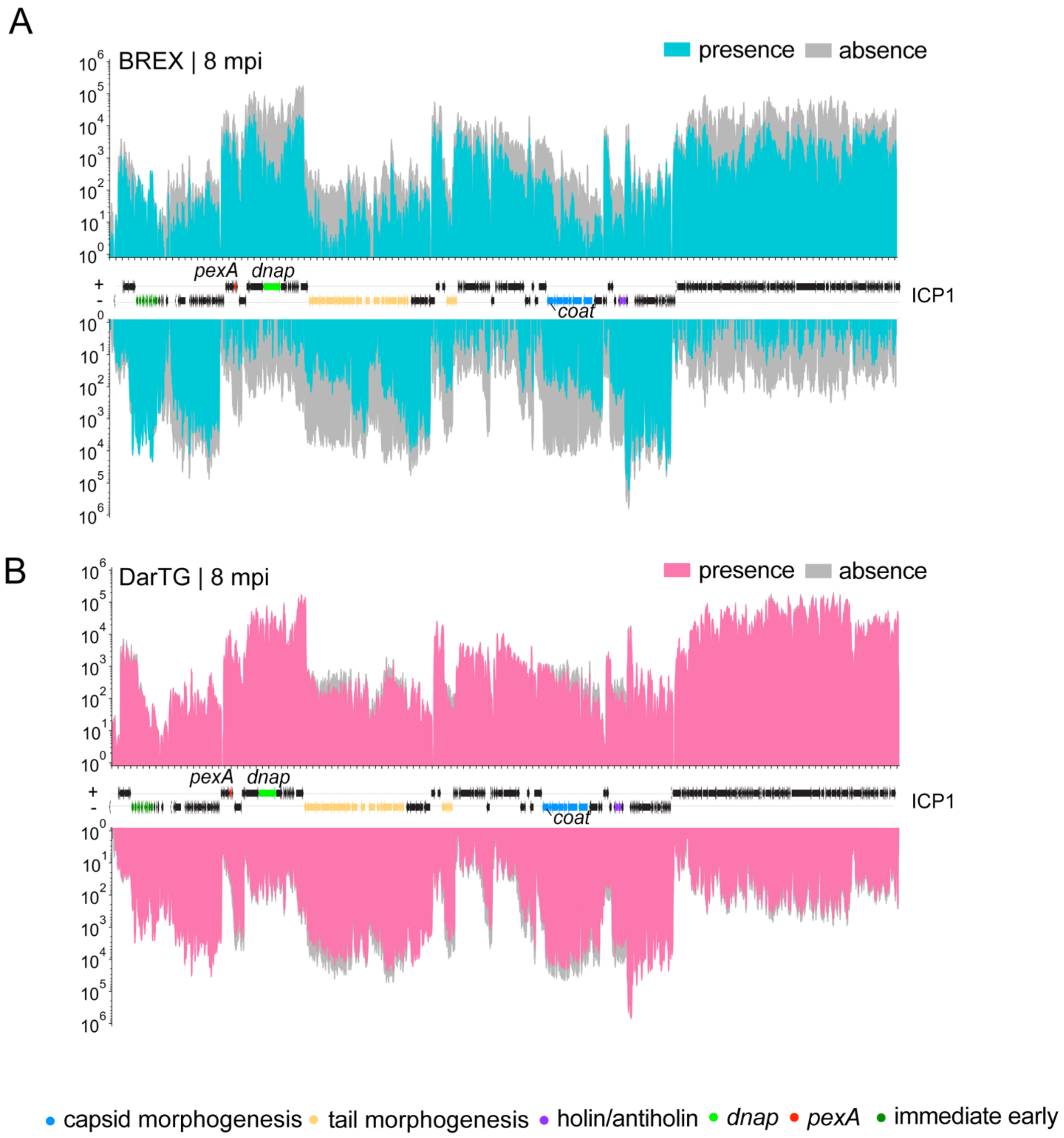


Fig. S2: (A, B) Average RNA-seq read coverage mapped to the ICP1 genome at 8 mpi in the presence or absence of (A) BREX and (B) DarTG. Forward strand (+) reads are plotted above the x-axis and reverse strand (–) reads below the x-axis. Coverage represents the mean of three biological replicates and is displayed on a log-scaled y-axis. ICP1 gene features are colored based on known or predicted functions according to the legend below panel B, while genes without functional annotation are shown in black.

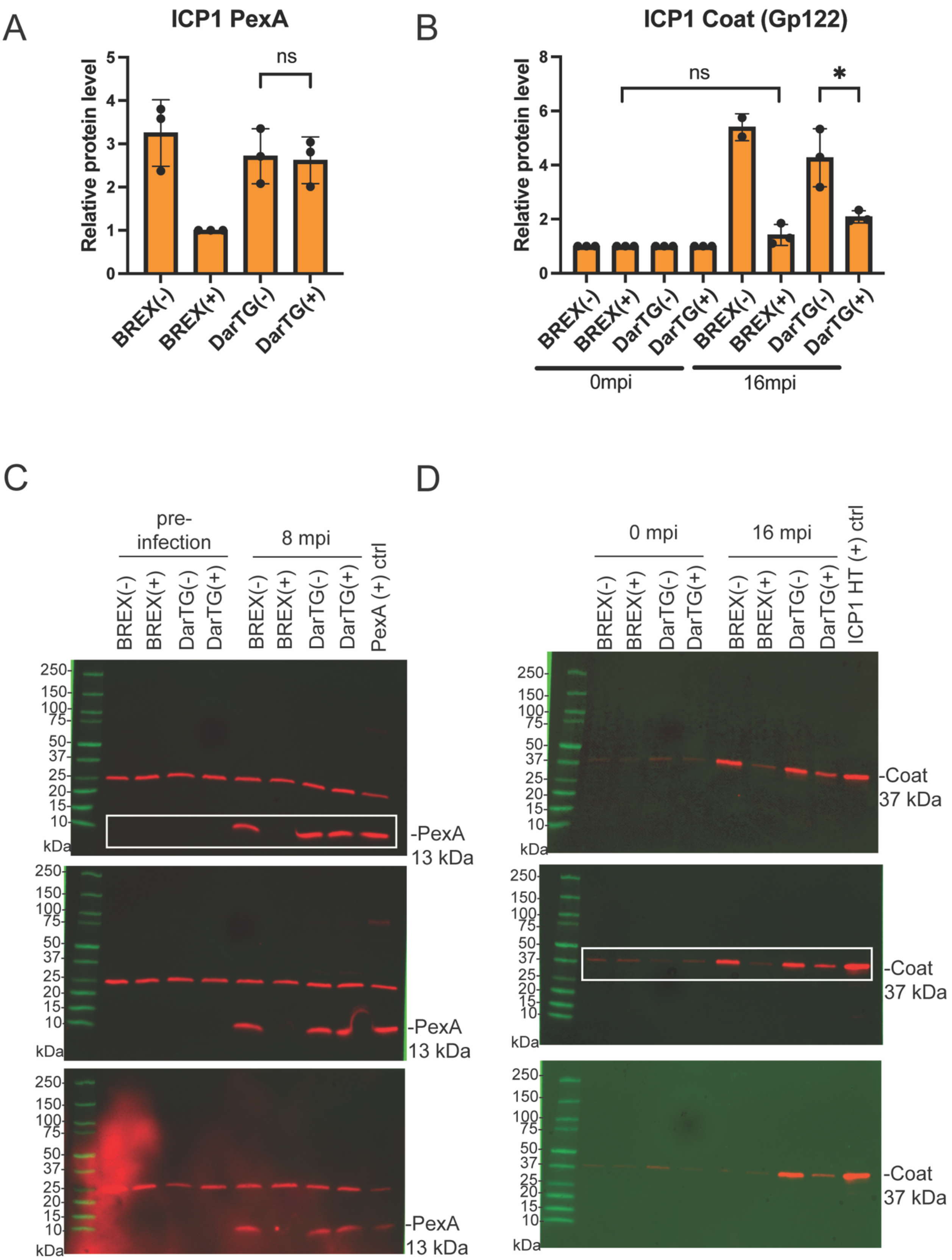

A
ICP1 PexA
Relative protein level
ns
BREX(-)
BREX(+)
DarTG(-)
DarTG(+)
B
ICP1 Coat (Gp122)
Relative protein level
ns
*
BREX(-)
BREX(+)
DarTG(-)
DarTG(+)
0mpi
16mpi
C
pre-infection
8 mpi
PexA (+) ctrl
kDa
-PexA 13 kDa
D
0 mpi
16 mpi
ICP1 HT (+) ctrl
kDa
-Coat 37 kDa

Fig. S3: (A, B) Relative abundance of ICP1-encoded (A) PexA (early) at 8 mpi and (B) Coat (late) at 16 mpi, respectively. Each dot represents an independent biological replicate, and bar heights represent the mean. Error bars represent standard deviation. Statistical significance was determined using an unpaired t-test (ns, not significant; *, $p < 0.05$). Protein levels were quantified from western blots and normalized to total protein using stain-free gel imaging (see methods). Because the coat protein is a structural component of the physical virion, normalization was performed against the corresponding phage inoculum (0 mpi) sample to accurately distinguish *de novo* protein synthesis from input phage. (C, D) Uncropped images for three biological replicates of the western blots for (C) PexA and (D) Coat from cells with and without BREX or DarTG before and after infection. The blots shown in the main text are highlighted with a white box. HT: High Titer stock of ICP1.

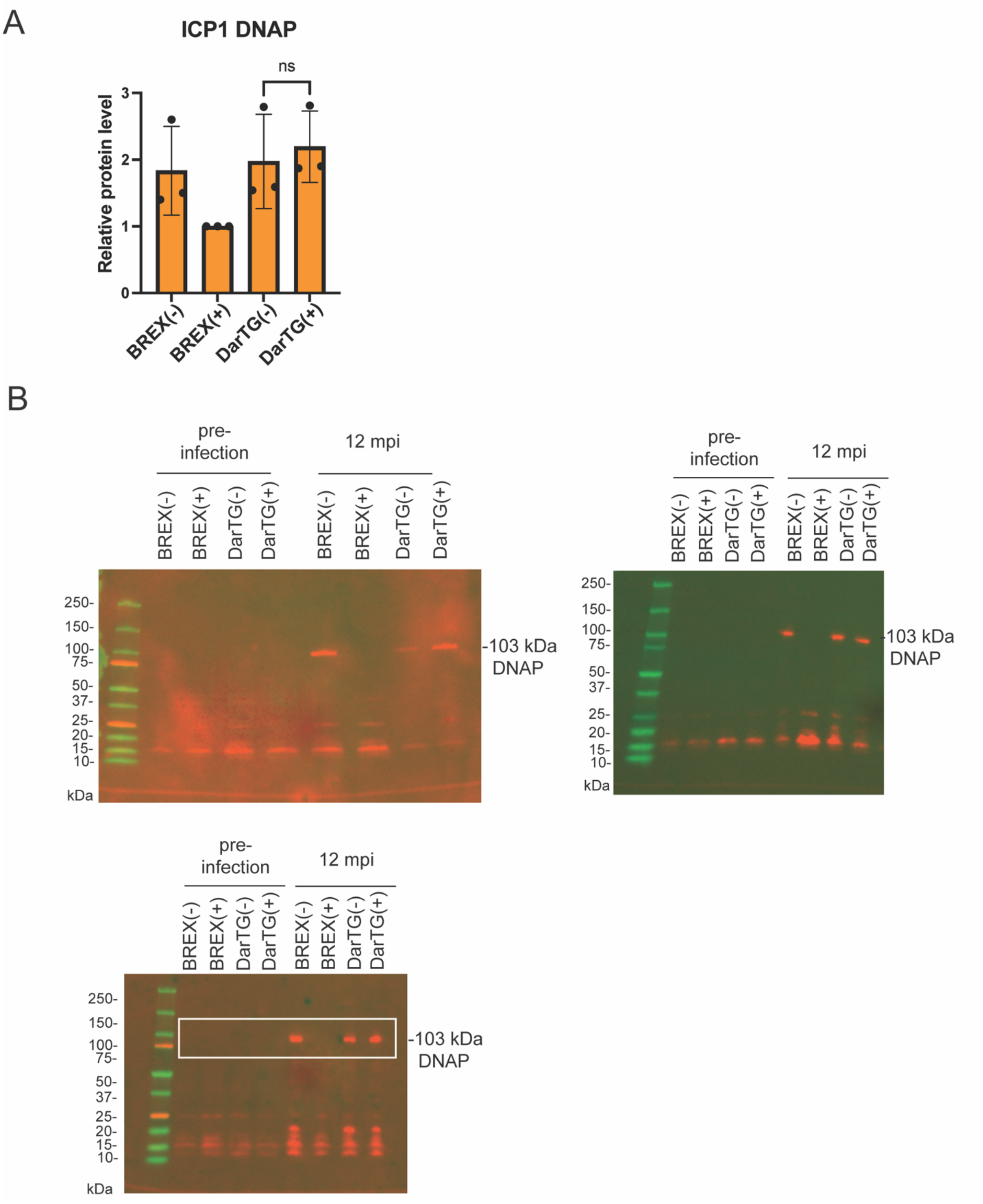
A
ICP1 DNAP
ns
Relative protein level
0
1
2
3
BREX(-)
BREX(+)
DarTG(-)
DarTG(+)
B
pre-infection
12 mpi
BREX(-)
BREX(+)
DarTG(-)
DarTG(+)
250-
150-
100-
75-
50-
37-
25-
20-
15-
10-
kDa
-103 kDa DNAP

Fig. S4: (A) Relative abundance of ICP1 encoded DNA polymerase (DNAP) at 12 mpi. Each dot represents an independent replicate, and bar heights indicate the mean. Error bars represent standard deviation. Statistical significance was determined using an unpaired t-test (ns, not significant; $p < 0.05$ was considered significant). Protein levels were quantified from western blots and normalized to total protein using stain-free gel imaging (see methods). (B) Uncropped images of three biological replicates of the western blots for DNAP from cells with and without BREX or DarTG before and 12 mpi. The blot shown in the main text is highlighted with a white box.

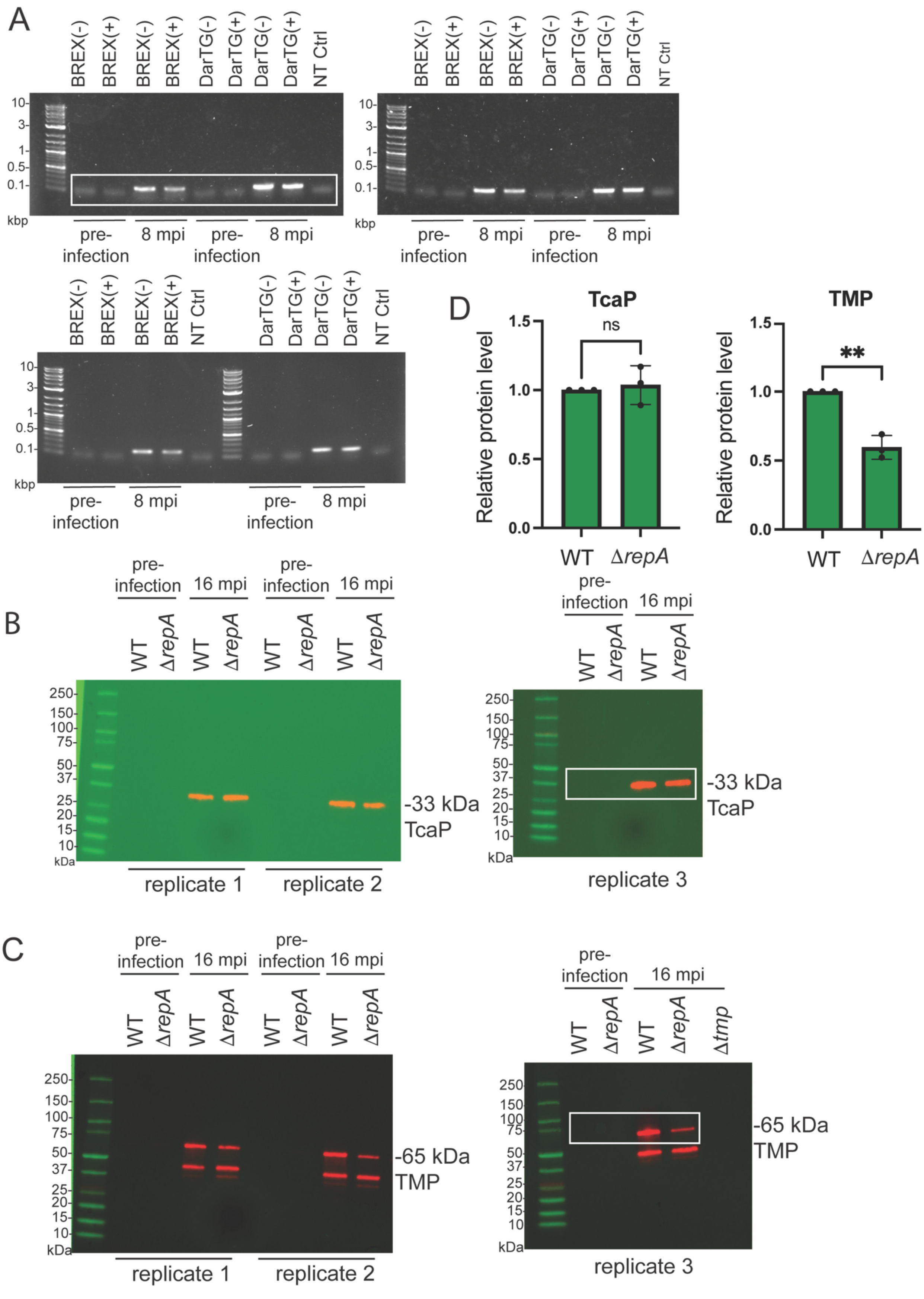
A
BREX(-) BREX(+) BREX(-) BREX(+) DarTG(-) DarTG(+) DarTG(-) DarTG(+) NT Ctrl
10- 3- 1- 0.5- 0.1-
kbp
pre-infection 8 mpi pre-infection 8 mpi
BREX(-) BREX(+) BREX(-) BREX(+) NT Ctrl DarTG(-) DarTG(+) DarTG(-) DarTG(+) NT Ctrl
D
TcaP
TMP
ns
**
Relative protein level
1.5 1.0 0.5 0.0
WT ΔrepA
B
pre-infection 16 mpi pre-infection 16 mpi
WT ΔrepA WT ΔrepA WT ΔrepA WT ΔrepA
250 150 100 75 50 37 25 20 15 10
kDa
-33 kDa TcaP
replicate 1 replicate 2 replicate 3
C
WT ΔrepA WT ΔrepA Δtmp
-65 kDa TMP

Fig. S5: (A) Three biological replicates of PCR analysis of PLE circularization before (t=0) and 8 mpi in the presence (+) or absence (−) of the indicated defense system. The gel shown in the main text is highlighted with a white box. L: Ladder, NT: No Template control. (B, C) Uncropped images for three biological replicates of the western blots of PLE encoded late proteins TcaP (B) and TMP (C) from cells with WT or ∆*repA* PLE before and 16 mpi by ICP1. (D) Relative abundance of TcaP and TMP at 16 mpi in *V. cholerae* with WT or ∆*repA* PLE. Each dot represents an independent replicate, and bar heights indicate the mean. Error bars represent standard deviation. Statistical significance was performed using an unpaired t-test (ns, not significant, **, $p < 0.005$). Protein levels were quantified from western blots and normalized to total protein using stain-free gel imaging (see methods). The blots shown in the main text are highlighted with a white box.

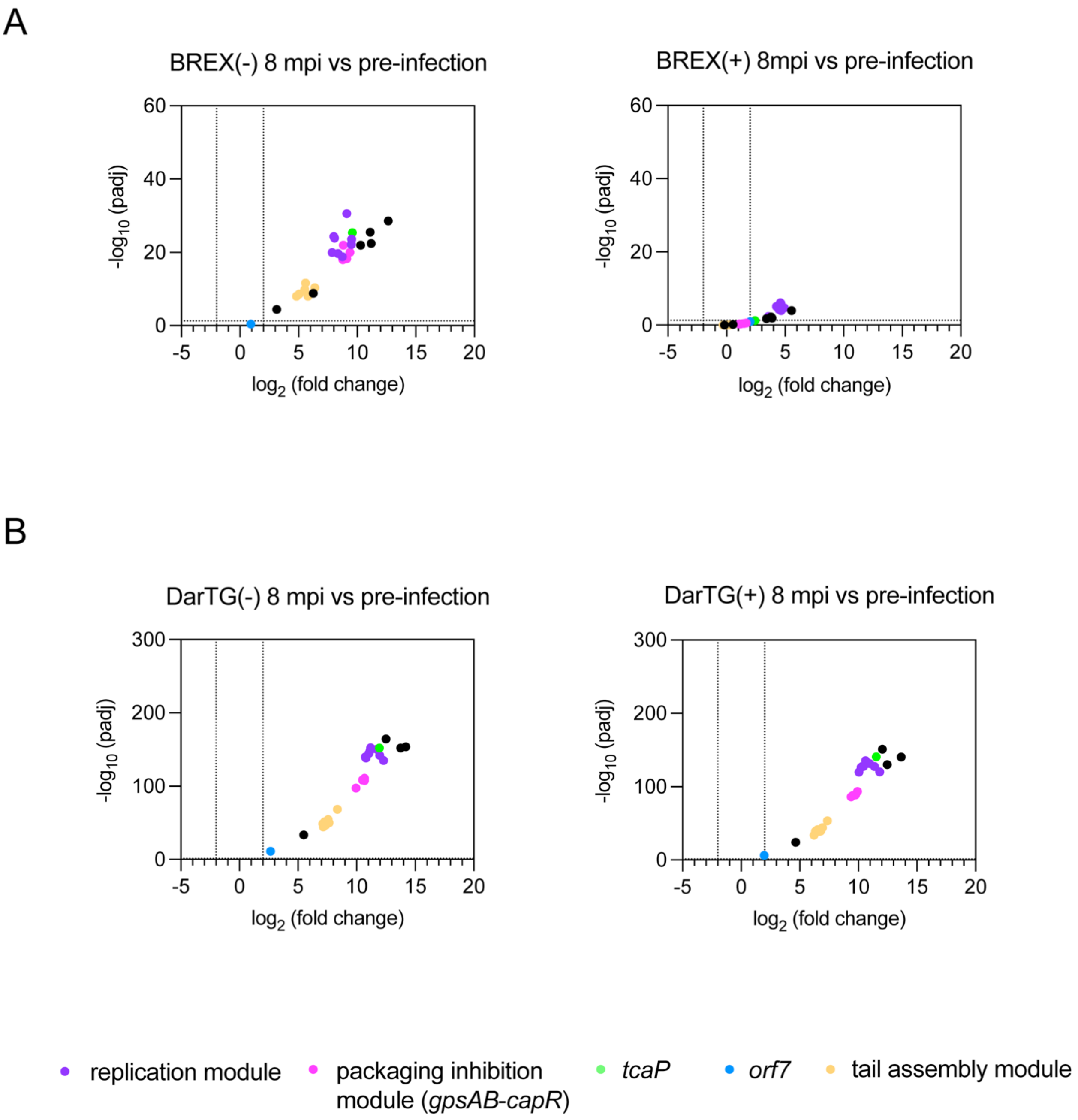


Fig. S6: (A, B) Volcano plot illustrating the differential expression of PLE genes comparing 8 mpi to pre-infection in the absence and presence of BREX (A) or DarTG (B). Vertical dotted lines indicate a $\log_2$ fold change cutoff of ≥ ±2, and horizontal dotted lines indicate a significance threshold of $-\log_{10} (p_{adj}) \geq 1.3$.

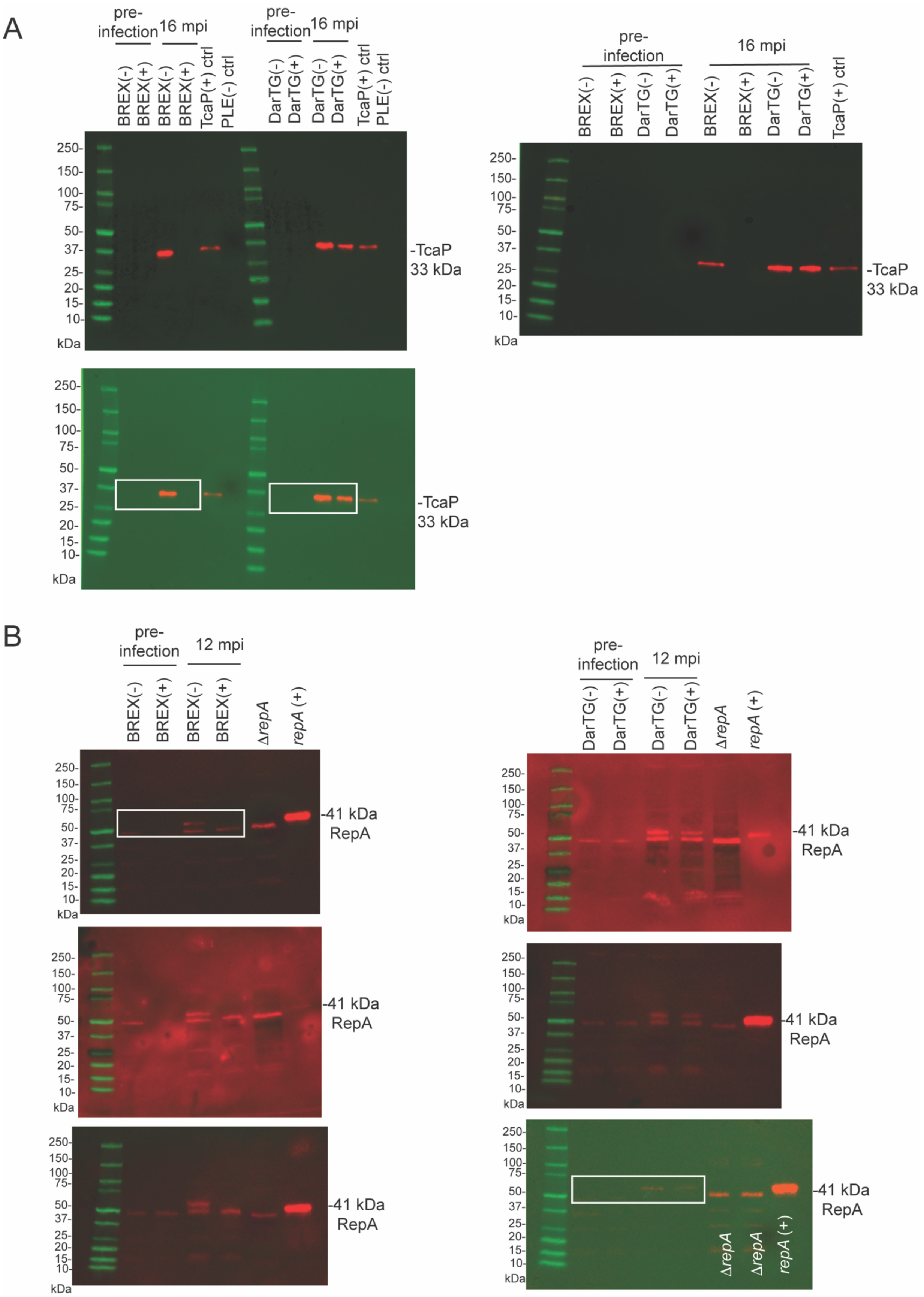

A
pre-infection
16 mpi
BREX(-)
BREX(+)
TcaP(+) ctrl
PLE(-) ctrl
DarTG(-)
DarTG(+)
250-
150-
100-
75-
50-
37-
25-
20-
15-
10-
kDa
-TcaP
33 kDa
B
12 mpi
ΔrepA
repA (+)
-41 kDa
RepA

Fig. S7: (A, B) Uncropped images of three biological replicates of the western blots for PLE-encoded late protein TcaP (A) and early protein RepA (B) at 16 mpi and 12 mpi, respectively. (B) Δ*repA* control is included to show that there was a band close to the expected size for RepA. The *tcaP* (+) and *repA* (+) controls are plasmid-based expression. See Table S14 for strain description. The blots shown in the main text are highlighted with a white box.

TableS14

| Strain ID | Designation | Description | Reference |
|---|---|---|---|
| TB354 | WT | *V. cholerae* E7946 PLE1 Δ*lacZ*::SpecR | This study |
| TB356 | BREX(-) | *V. cholerae* E7946 PLE1 Δ*lacZ::*SpecR *Vch*Ind5$^+$ Δhotspot5 | This study |
| TB359 | BREX(+) | *V. cholerae* E7946 PLE1 Δ*lacZ*::SpecR *Vch*Ind5$^+$ | This study |
| TB458 | DarTG(-) | *V. cholerae* E7946 PLE1::KanR PDE$^+$ Δ*old*::frt Δ*darT*::frt Δ*lacZ*::SpecR | This study |
| KMP168 | DarTG(+) | *V. cholerae* E7946 PLE1::KanR PDE$^+$ Δ*lacZ*::SpecR | [23] |
| CMB555 | *tcaP* + ctrl | *E.coli* BL21 pETDUET P$_{tac}$-*gp122*::6xhis, P$_{tac}$-*tcaP* | [15] |
| MR151 | *tmp* - ctrl | *V. cholerae* E7946 PLE1::KanR Δ*tmp* (*orf22*)::frt-SpecR | This study |
| ACM451 | *pexA* + ctrl | *V. cholerae* E7946 PLE1::KanR containing pKL06.2 *pexA* (*gp51)* | [13] |
| TB362 | Δ*repA* | *V. cholerae* E7946 PLE1 Δ*repA*::frt Δ*lacZ*::SpecR | This study |
| KS1828 | *repA* + ctrl | *V. cholerae* E7946 pKL06.2 *repA* | This study |
| TBphi1 | ICP1 phage | ICP1_2006_Dha_E (Accession MH310934) ΔCRISPR Δ*cas2-3* Δ*orbA* | This study |
